# A Multiscale Computational Architecture to Study Signaling Dynamics at Cell-Cell Interfaces

**DOI:** 10.64898/2026.03.16.712104

**Authors:** Yinghao Wu

## Abstract

Intercellular communication is governed by the spatiotemporal dynamics of protein complexes at the cell-cell interface. However, conventional static interaction models fail to incorporate key physical constraints, such as steric hindrance, spatial compartmentalization, and dimensionality reduction that regulate complex assembly in vivo. To bridge the gap between static network topology and dynamic systems biology, we developed a multi-scale computational framework. We first identified a highly conserved, Fibroblast Growth Factor Receptor 1 (FGFR1)-centered cell adhesion and signaling motif by analyzing a diverse set of human cell–cell interfaces. We then constructed a multi-layer spatial stochastic simulator to recapitulate and interrogate the dynamic behavior of this network motif at cell-cell interfaces. Atomic-resolution structural models of the protein complexes within the motif were further generated using AlphaFold to define interaction rules for the stochastic simulations by categorizing binding interfaces. Our results show that the structural arrangement of cell-cell adhesion complexes controls how FGFR1 receptors cluster at the cell–cell interface, effectively dividing the membrane into distinct functional microdomains. Competition from decoy receptors further regulates this process by capturing receptors before they can participate in signaling. Even small changes in binding affinity can therefore alter receptor organization and disrupt normal signal transduction, which may contribute to human disease. By integrating macro-scale interactomics, atomic-level structural bioinformatics, and mesoscale stochastic modeling, this study reveals how structural interaction rules, combined with spatial constraints, shape the formation and function of intercellular signaling networks.

## Introduction

Intercellular communication is a fundamental driver of multicellular biology [1, 2] governing a wide range of physiological processes from embryonic development [3] to tissue homeostasis [4]. This communication is tightly regulated by the dynamic interplay between cell–cell adhesion, which establishes the physical and mechanical architecture of cellular boundaries, and cell signaling, which converts extracellular cues into precise biochemical responses [5]. Rather than functioning independently, adhesion and signaling are deeply interconnected: adhesive complexes spatially organize signaling receptors [6], while localized signaling events continuously remodel the adhesive landscape [7]. Consequently, disruptions in the coordination between these two processes are a major contributor to numerous pathologies, including cancer metastasis, autoimmune disorders, and neurodevelopmental diseases [8]. Despite its central biological importance, the biophysical and structural mechanisms that govern the spatiotemporal regulation of cell–cell communication remain incompletely understood. Elucidating these dynamic mechanisms is therefore of significant importance, as it will not only advance our fundamental understanding of complex physiological activities, but also provide critical insights for identifying novel and highly specific therapeutic targets aimed at modulating aberrant intercellular junctions [9].

Traditionally, the investigation of complex intercellular signaling networks has relied on a combination of classical biochemistry, structural biology, and advanced fluorescence microscopy. Biochemical techniques such as co-immunoprecipitation are invaluable for identifying interacting protein partners [10, 11]; however, these bulk population assays inherently disrupt the delicate spatial architecture of cellular junctions, obscuring the localized dynamics of protein complex assembly. Conversely, structural methods such as X-ray crystallography [12–16] and cryo-electron microscopy [17–19] provide exquisite atomic-resolution snapshots of isolated protein complexes, yet they remain fundamentally static and cannot capture the temporal, multi-protein competition and coordination that occurs within a living membrane environment. Although live-cell imaging and super-resolution microscopy [20–24] have significantly improved our ability to track molecular movements in real time, interrogating the narrow and densely packed intercellular cleft remains technically challenging. These optical approaches often lack the resolution necessary to distinguish competing structural interfaces in vivo and are further limited by the perturbative effects of fluorescent labeling. Consequently, current experimental paradigms struggle to simultaneously integrate the physical constraints of the cell–cell boundary, such as dimensionality reduction, and spatial compartmentalization, with the kinetic rates of molecular interactions. This methodological gap prevents a fully quantitative and predictive understanding of how atomic-scale binding rules scale up to govern dynamic, mesoscale communication networks.

To overcome the limitations of traditional experimental approaches, computational methods have become increasingly important for investigating intercellular communication. Numerous online databases and network analysis tools now enable high-throughput mining of protein–protein interactions (PPIs) [25]. However, these resources typically generate static, topological maps of interaction networks and lack information about the physical properties and geometric constraints of the underlying binding interfaces. Structural bioinformatics [26–28], particularly recent advances driven by AlphaFold [29], can resolve these interactions at atomic resolution, but such structural models remain inherently static and do not capture the temporal dynamics or competitive assembly processes of multi-protein complexes. Molecular dynamics (MD) simulations can provide detailed insights into these dynamic behaviors [30–37], yet they are computationally prohibitive for modeling large multi-protein systems at cell–cell interfaces over biologically relevant timescales. In contrast, mesoscale stochastic simulations can capture subcellular dynamics with appropriate spatial and temporal resolution over seconds to minutes [38–40]. However, the predictive power of these models depends critically on the accuracy of the predefined reaction rules and physical constraints used as inputs. Because each computational approach has distinct strengths and limitations, integrating them into a unified multiscale framework represents a powerful strategy for bridging the gap between static interactomic maps and dynamic systems-level understanding of intercellular communication.

To implement this strategy, we developed a multi-scale computational framework to study the dynamic behavior of cell-cell interfaces. We first analyzed existing databases to identify a highly conserved signaling network centered around the Fibroblast Growth Factor Receptor 1 (FGFR1). Next, we used AlphaFold to map the structural interfaces of these proteins, giving us clear physical rules for how they bind and compete. We then fed these structural rules into a spatial stochastic simulator. By explicitly dividing the biological environment into discrete 3D spatial compartments, such as the extracellular area, the plasma membrane, and the cytoplasm, this simulator accurately captures how molecules move and react within restricted physical boundaries. Our results show that different adhesion molecules, such as NECTIN1 and L1CAM, physically organize the membrane to control where and how efficiently FGFR1 signals. Furthermore, we demonstrate that decoy receptors (FGFRL1) act as a critical “gatekeeper” to tune the overall signal strength, a delicate balance that is disrupted by a specific gene mutation linked to Kallmann syndrome. Ultimately, this study highlights how combining structural biology with spatial compartmentalization can reveal the hidden physical rules that govern intercellular communication.

## Methods

### Overview of the computational framework

Our multi-scale methodological framework (**Figure 1**) provides a rigorous, vertically integrated pipeline that bridges bioinformatics discovery, atomic-scale structural proteomics, and mesoscale spatial stochastic modeling to elucidate the biophysical principles governing membrane-bound signaling hubs. The process is initiated by a high-throughput Database Search across curated repositories, including CITEdb [41], Cellinker [42], and the Human Protein Atlas [43], to identify a Conserved Network Motif. Following motif identification, the framework bifurcates into two essential, parallel analytical tracks that define the fundamental physics of the system. In the Structural Analysis track, we utilize state-of-the-art AlphaFold3 predictions to map the precise atomic-scale binding interfaces of the network components, allowing us to categorize whether two protein-protein interactions can either coexist through their non-overlapping binding interfaces or compete against each other due to the overlapping interfaces. Simultaneously, the Cell Environment panel establishes the constraints for molecular motion and boundaries of biological interface.

**Figure 1:**
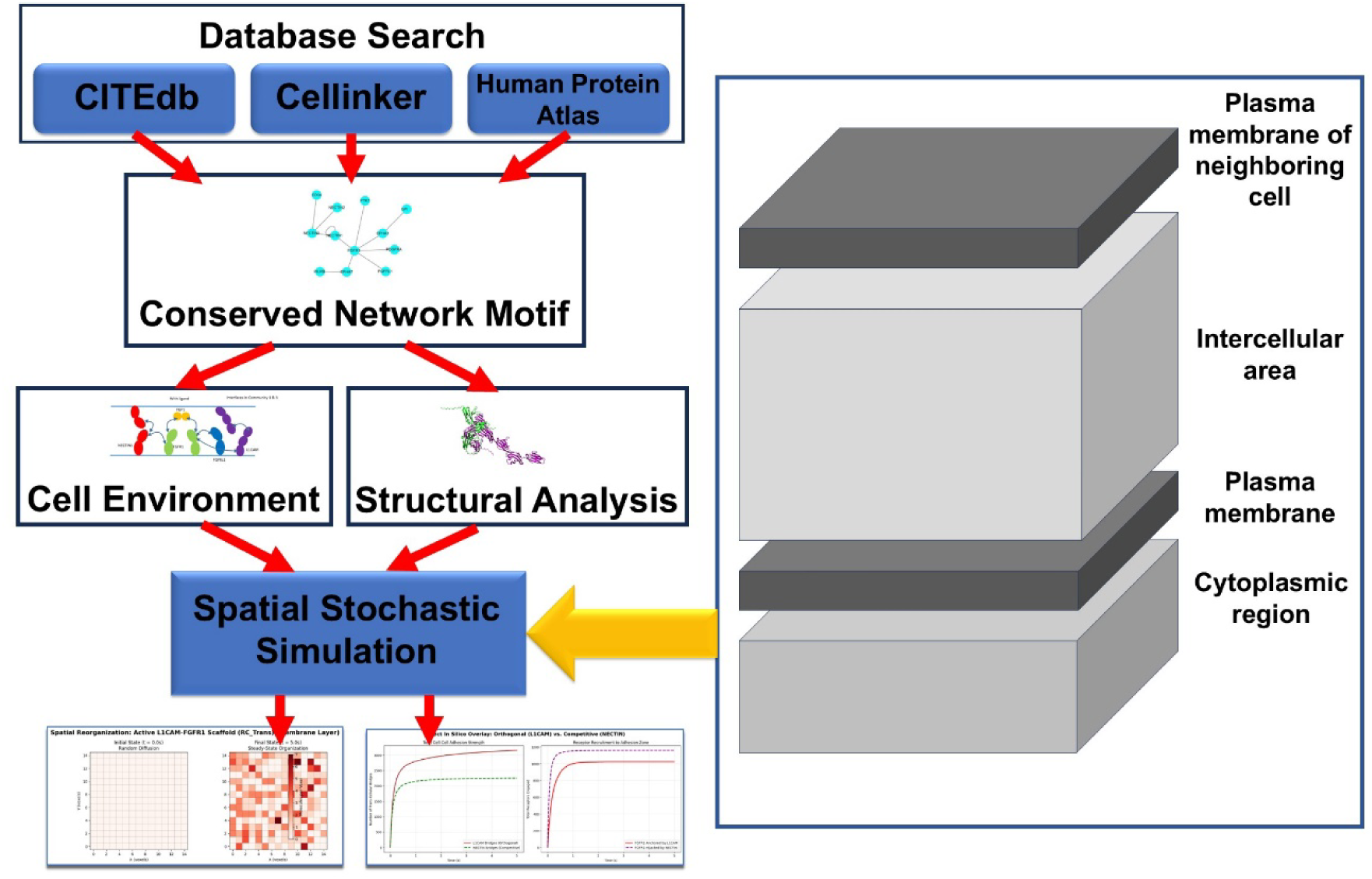
Our multiscale computational framework integrate network analysis of protein-protein interactions, atomic-level structural bioinformatics, and mesoscale spatial stochastic simulations.

These two elements are then synthesized within a Spatial Stochastic Simulation utilizing a Reaction-Diffusion Master Equation (RDME) engine [44–46] implemented via a discretized Gillespie Algorithm [47], which models the cell-cell interface as a 15×15×5 voxel grid with a characteristic scale of 0.3μm. This architecture utilizes a functional Z-axis to compartmentalize the biological environment into a multi-layer geometry, separating the plasma membrane of neighboring cell, the intercellular cleft, and the plasma membrane of target cell and its cytoplasmic regions. The simulation progresses by stochastically sampling two discrete event types: Reaction Events, where molecular transformations occur within a single local sub-volume based on binding propensities, and Diffusion Events, where molecules “jump” between adjacent voxels at rates defined by *D/h^2^*, where *h=0.3*μ*m* is the voxel length and *D* is the diffusion constant. The iterative loop stochastically advances the simulation by summing the propensities of all possible reactions and diffusion jumps to calculate a random waiting time between discrete molecular events. A single event is then selected based on its relative probability and executed, updating the voxel-level molecular counts before the cycle restarts.

### Discovery of the conserved network motif

To identify biologically conserved adhesion and signaling modules, we developed a multi-database bioinformatics pipeline that systematically filters established protein–protein interactions (PPIs) by their spatial and contextual relevance. We began by mining CITEdb, which catalogs human cell–cell interactions across diverse physiological and pathological states [41]. From this repository, we extracted 41 unique cell–cell interfaces, representing critical functional boundaries such as those between macrophages and endothelial cells. In parallel, we queried the Cellinker database to compile a comprehensive network of intercellular communication [42]. To ensure our framework modeled true mechanical junctions rather than transient paracrine signaling, we rigorously filtered this dataset to exclude non-physical contacts, yielding a curated set of 1,744 structural PPIs localized exclusively to the plasma membrane.

Recognizing that the mere structural capacity to bind does not guarantee in vivo interaction, we cross-referenced these candidate pairs with tissue-specific data from the Human Protein Atlas [43]. Because protein expression varies drastically across cell lineages, a candidate PPI was only retained for a given interface if both constituent proteins exhibited medium-to-high expression levels in their respective apposing cell types. By aggressively filtering out undetectable or low-expressing proteins, we isolated interaction networks that are biologically plausible at specific cellular junctions. Ultimately, analyzing these validated networks revealed a distinct subset of PPIs highly conserved across major cell–cell interface communities. These conserved interactions naturally organize into recurrent, modular network motifs that are reused across diverse cellular environments. Most notably, this analysis uncovered a dominant core signaling module centered at FGFR1 (**Figure 2a**) [48], highlighting its ubiquitous cross-talk with various cell adhesion molecules at the cell-cell interfaces, including NECTIN1 [49] and L1CAM [50].

**Figure 2:**
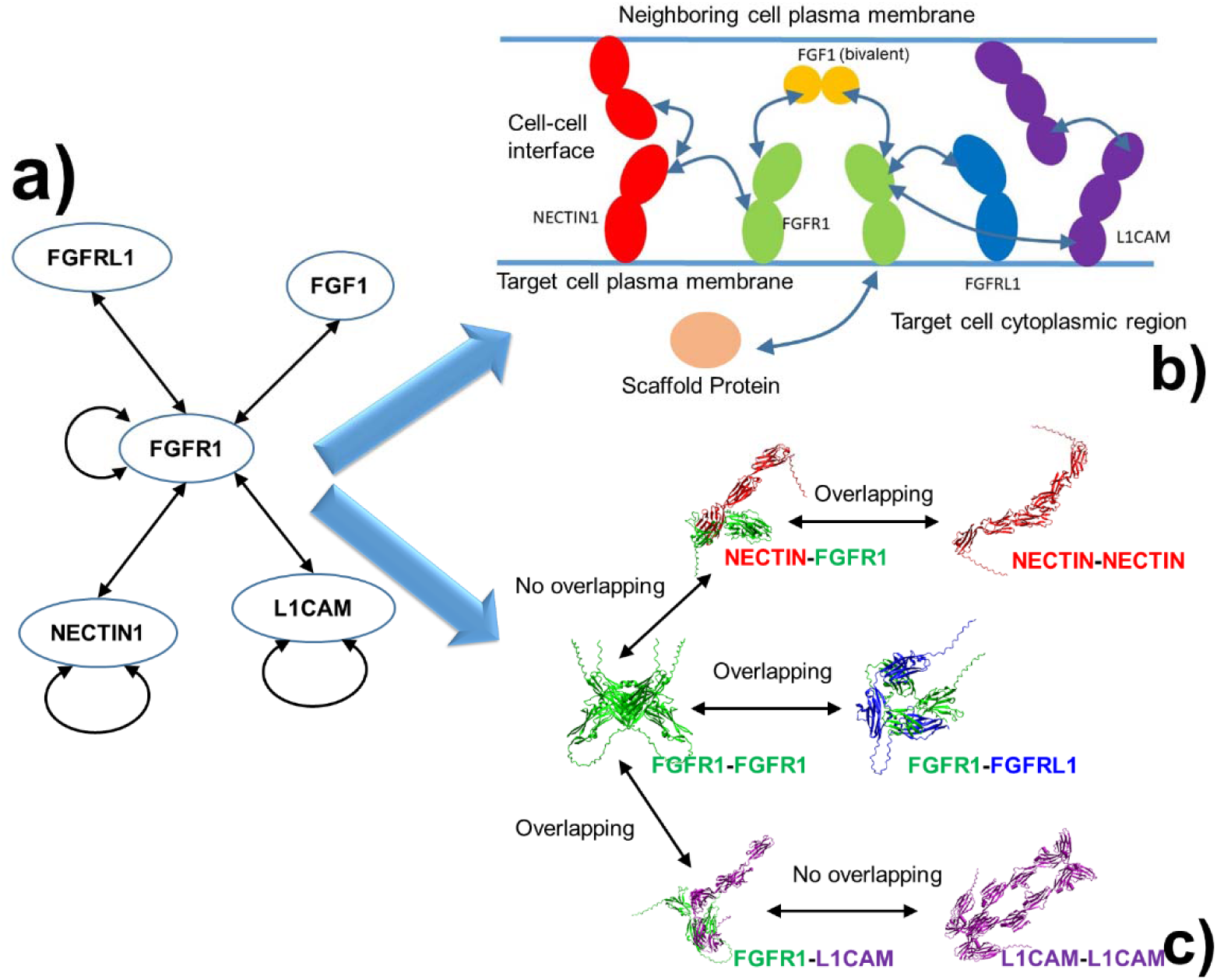
We identified a conserved cell adhesion and signaling motif centered at the Fibroblast Growth Factor Receptor 1 (FGFR1) (a). We transformed the motif into a physical architecture with the cellular context of the cell-cell interface (b). In parallel, we generated a set of reaction rules for protein-protein interactions in the motif based on the structural analysis (c).

### Setup of spatial environments and reaction rules

Given the network motif of PPIs, we transformed it into a physical architecture with the cellular context of the cell-cell interface. As shown in **Figure 2b**, the top layer represents the plasma membrane of the neighboring cell, populated by the trans-cellular binding partners (e.g., the top-cell fractions of NECTIN1 and L1CAM). Molecular diffusion in this layer is restricted to 2D lateral movement. The intercellular area separates the two cells. This is the compartment where 3D diffusion is permitted, allowing the soluble FGF1 ligand to move freely between two cell surfaces. The primary signaling surface of the target cell. This 2D plane contains the full suite of signaling receptors (FGFR1), decoy receptors (FGFRL1) [51], and the bottom-cell fractions of the cell adhesion molecules. Below the plasma membrane of target cell, we provide the necessary volumetric depth to represent the intracellular space where scaffold proteins can move freely in 3D.

After defining the spatial environment of cell-cell interface, we further generated a set of reaction rules for PPIs in the motif based on the structural analysis. Specifically, we analyzed the overlap of protein–protein binding interfaces among several FGFR1-centered interactions. For each PPI, we first generated structural models of the corresponding protein complexes using AlphaFold3. Binding interfaces were then defined by extracting residues whose side-chain atoms were within 3 Å of any atom from the interacting partner in the predicted complex. Interface overlap between two PPIs was assessed by comparing the sets of residues involved in binding, allowing us to determine whether distinct partners compete for the same structural surface or if they can assemble simultaneously. These atomic-scale structural geometries were directly translated into the biochemical reaction rules governing our spatial stochastic simulation. By distinguishing between mutually exclusive and co-existing interfaces, we established three primary rule sets that dictate the mesoscale behavior of the network (**Figure 2c**).

First, the *cis*-interaction between NECTIN1 and FGFR1 overlaps with its *trans*-adhesion interface, forcing a mutually exclusive choice between cell-cell bridging and receptor binding. However, because the FGFR1 dimerization site remains accessible to NECTIN1, NECTIN-anchored receptors can still cross-link into active signaling complexes. Second, the decoy receptor FGFRL1 physically occludes the canonical FGFR1 homo-dimerization interface, strictly preventing active dimer formation. Third, L1CAM utilizes an orthogonal scaffolding mechanism; while its *cis*-interaction with FGFR1 also blocks receptor dimerization, it does not interfere with L1CAM *trans*-adhesion. Consequently, L1CAM can simultaneously bridge neighboring cells and dock FGFR1 monomers in *cis*. Finally, scaffold proteins in the cytoplasm can bind to receptors through their intracellular domains, but only after the receptor are engaged with extracellular ligands and form ligand-receptor oligomers. All the reactions involved in the network motif are summarized in **Table 1**.

**Table 1:** List of all reactions of cell-cell adhesion and signaling involved in the simulation system.

| index | Reactant_1 | Reactant_2 | Product | Biological_Process |
| --- | --- | --- | --- | --- |
| 1 | Nectin (top cell) | Nectin (bottom cell) | Trans Nectin adhesion complex | Formation of trans-cell Nectin adhesion |
| 2 | Fibroblast Growth Factor Receptor 1 (FGFR1) | Nectin (bottom cell) | FGFR1-Nectin complex | Receptor capture by Nectin scaffold |
| 3 | L1 Cell Adhesion Molecule (L1CAM, top cell) | L1 Cell Adhesion Molecule (L1CAM, bottom cell) | Trans L1CAM adhesion complex | Formation of trans L1CAM adhesion |
| 4 | Fibroblast Growth Factor Receptor 1 (FGFR1) | L1CAM (bottom cell) | FGFR1-L1CAM complex | Receptor capture by L1CAM scaffold |
| 5 | Fibroblast Growth Factor Receptor 1 (FGFR1) | Trans L1CAM adhesion complex | FGFR1-Trans L1CAM complex | Receptor recruitment to L1CAM bridge |
| 6 | L1CAM (top cell) | FGFR1-L1CAM complex | Trans L1CAM-FGFR1 complex | Completion of L1CAM receptor scaffold |
| 7 | Fibroblast Growth Factor (FGF ligand) | Fibroblast Growth Factor Receptor 1 (FGFR1) | FGFL-FGFR1 complex | Ligand binding to receptor |
| 8 | Fibroblast Growth Factor (FGF ligand) | FGFR1 dimer | FGFL-FGFR1 dimer complex | Ligand binding to receptor dimer |
| 9 | Fibroblast Growth Factor (FGF ligand) | FGFR1-Nectin complex | FGFL-FGFR1-Nectin complex | Ligand binding to Nectin-associated receptor |
| 10 | Fibroblast Growth Factor (FGF ligand) | FGFR1 dimer-Nectin complex | FGFL-FGFR1 dimer-Nectin complex | Ligand binding to Nectin-associated receptor dimer |
| 11 | Fibroblast Growth Factor Receptor 1 (FGFR1) | Fibroblast Growth Factor Receptor 1 (FGFR1) | FGFR1 dimer | Receptor dimerization |
| 12 | Fibroblast Growth Factor Receptor 1 (FGFR1) | FGFR1 decoy receptor | FGFR1-Decoy complex | Receptor sequestration by decoy |
| 13 | FGFL-FGFR1 complex | Fibroblast Growth Factor Receptor 1 (FGFR1) | FGF-FGFR1 dimer (active signaling complex) | Ligand-induced receptor crosslinking |
| 14 | FGFL-FGFR1 complex | FGFR1-Nectin complex | FGFL-FGFR1 dimer-Nectin complex | Crosslinking on Nectin scaffold |
| 15 | FGF-FGFR1 dimer<br>(active signaling<br>complex) | Intracellular Scaffold<br>Protein | Ligand-Receptor-Scaffold signaling<br>complex | Cross-membrane<br>Signal Transduction |

### Determination of simulation parameters

With the spatial architecture and structural reaction rules established, the next critical step was to parameterize the dynamic behavior of the system by quantifying molecular mobility and interaction speeds.

The diffusion constants govern the jump rates between voxels in the Spatial Gillespie algorithm. We defined three distinct physical regimes based on established biophysical literature: 1) Soluble proteins (such as FGF1 and scaffold protein) diffusing in the aqueous extracellular cleft or cytoplasmic region move rapidly. 2) Monomeric proteins anchored in the lipid bilayer experience massive viscous drag compared to aqueous proteins. Typical transmembrane receptors (like FGFR1 or free NECTIN) diffuse roughly 10 to 100 times slower than free soluble ligands. 3) Once a receptor binds a ligand, forms a dimer, or tethers to other adhesive proteins, its effective hydrodynamic radius and physical tethering increase significantly, dropping its mobility. We scaled complex diffusion to be roughly 4 times slower than free monomers in the plasma membrane.

In terms of reactions, because exact, context-specific *k_on_*and *k_off_* rates for every possible complex at the living cell-cell interface are impossible to measure with current experimental technology, the strategy of relative kinetic hierarchies was applied. Specifically, we scaled the binding probabilities to reflect the known biophysics of the system. First, the binding of a soluble ligand to a free monomeric membrane receptor is highly efficient. Once the fast-moving ligand enters the voxel containing a receptor, binding is highly probable. Moreover, the binding between a receptor-bound ligand to another receptor to form a ligand-receptor oligomer is even faster due to membrane confinement. Second, the homophilic *trans-*interactions of cell adhesion molecules such as L1CAM NECTIN1 are relatively faster to physically anchor the cells together than the thermal noise that pushes cells apart. Third, *cis*-interactions between membrane receptors from the same cell are slower than the *trans*-interactions because they requires highly specific orientation and alignment. Finally, to translate these macroscopic relative rates into probabilities for the Gillespie engine, we applied the standard Reaction-Diffusion Master Equation (RDME) volume scaling. Detailed values of all diffusion and reaction parameters used in the simulation can be found in **Table 2**.

**Table 2:**
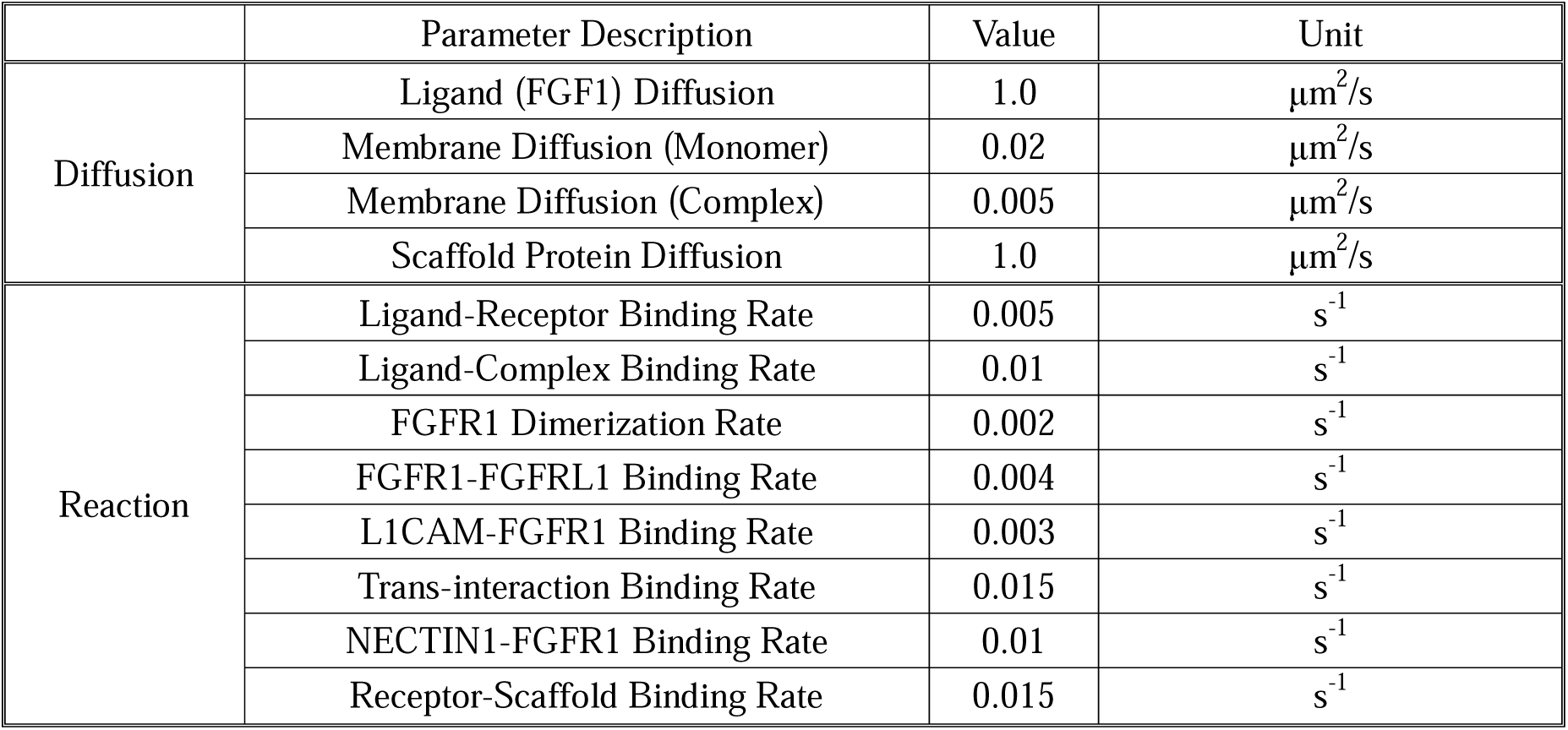
Parameters of diffusion and reaction rates and their corresponding values adopted in the simulations.

### Code availability

All relevant source codes of the CNM classification and Interface Calculation can be found in the GitHub repository: https://github.com/wujah/SSS_CCI

## Results

We first simulate the process of cross-membrane signal transduction without the impact of decoy receptor FGFRL1 and the crosstalk with other cell adhesion molecules. Simulation results are shown in **Figure 3a**. The curves in the figure reveal a sequential assembly process. Free ligand rapidly decreases (blue) as it binds membrane receptors to form the single bound ligand-receptor complex. This transient complex subsequently recruits additional receptors or receptor dimers to form cross-linked complexes. As a result, the pool of free receptors declines quickly, while single bound ligand-receptor complex rises and reaches a steady-state level (purple), reflecting a dynamic balance between formation and consumption. On the other hand, the cross-linked complex formed between bivalent ligands and dimeric receptors exhibits a transient peak before decreasing (green), indicating that it functions as an intermediate species. This decline corresponds to the recruitment of intracellular scaffold proteins, which bind the bivalent ligands and dimeric receptors complex to form the final active signaling hub. Consistent with this mechanism, the number of free scaffold proteins gradually decreases (cyan) while the signaling hubs accumulates monotonically over time (black).

**Figure 3:**
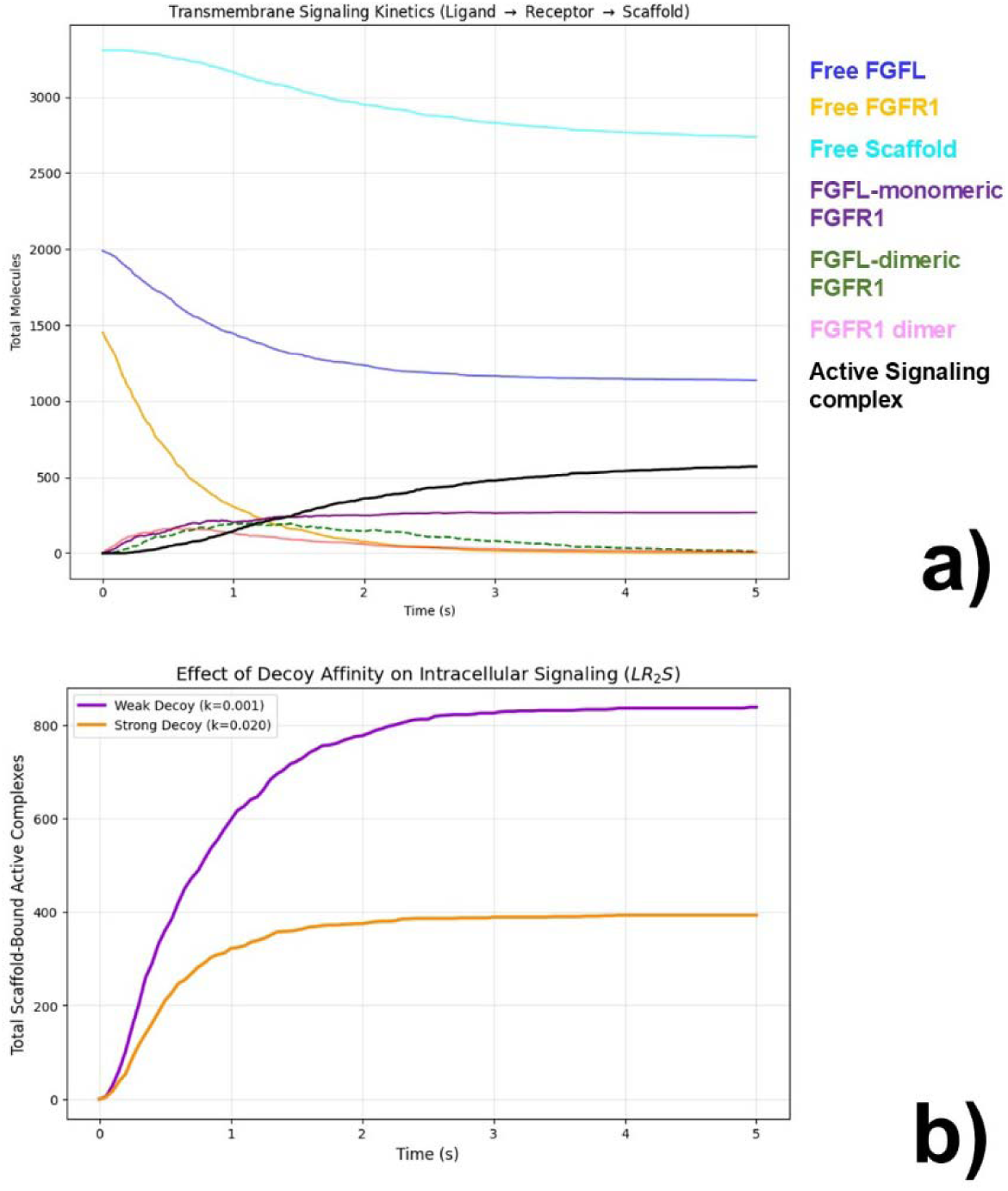
We first simulate the process of cross-membrane signal transduction without the impact of decoy receptor FGFRL1 and the crosstalk with other cell adhesion molecules. Our results highlight how multivalent ligand binding, receptor cross-linking, and scaffold recruitment cooperate to generate robust transmembrane signaling dynamics **(a)**. We further include decoy receptor FGFRL1 into the system which can compete with receptor dimerization. We show that by increasing the stability of the FGFR1-FGFRL1 complex, receptors are sequestered into inactive states more efficiently **(b)**.

Notably, the formation of the active signaling hub occurs much more slowly. Although the ligand-receptor complexes accumulate rapidly at early times, the cross-membrane signal transduction requires the recruitment of intracellular scaffold proteins from the cytoplasm to the membrane. This additional step introduces both spatial and kinetic constraints, including scaffold diffusion from the cytoplasm and the requirement for productive encounters with membrane-bound ligand-receptor complexes. This delayed accumulation of signaling hub highlights an important biological principle: signal activation is not determined solely by ligand binding, but by the assembly of higher-order receptor–scaffold complexes that stabilize signaling at the membrane. The requirement for scaffold recruitment effectively acts as a regulatory checkpoint, ensuring that transient receptor engagements are filtered before being converted into sustained intracellular signaling. Together, these results highlight how multivalent ligand binding, receptor cross-linking, and scaffold recruitment cooperate to generate robust transmembrane signaling dynamics.

We further include decoy receptor FGFRL1 into the system which can compete with receptor dimerization. To understand how the decoy receptor regulates the system, we compared two different binding rates between the receptor and the decoy. In the first scenario, the binding rate is low. In this case, receptor monomers are more likely to collide with each other or bind with ligands to form active dimers. This leads to a steady increase in the concentration of scaffold-bound active complexes over the 5-second simulation period (purple curve in **Figure 3b**). In the second scenario, the binding rate between the receptor and the decoy is high. Under these conditions, the decoy outcompetes the signaling pathway for the available receptor monomers. Instead of forming active dimers, the receptors are rapidly pulled into inactive receptor-decoy complexes. Consequently, the final amount of intracellular signaling is significantly lower than in the first scenario (orange curve in **Figure 3b**).

This kinetic competition provides a physical framework for understanding the D129A variant of the FGFR1 gene, which is linked to Kallmann syndrome [52]. AlphaFold model shows that the D129 residue is located at the binding interface between FGFR1 and its partner. Binding affinity calculations performed with PRODIGY [53, 54] further indicates that the D129A substitution strengthens the interaction between these two proteins from –18.8kcal/mol to – 19.6kcal/mol. By increasing the stability of the FGFR1-FGFRL1 complex, the mutation ensures that receptors are sequestered into inactive states more efficiently. This disruption of normal signaling dynamics likely explains the impaired development of hormone-producing neurons observed in patients.

To understand how different cell adhesion molecules influence signaling, we compared two distinct systems: The first one is only driven by NECTIN1 and the second one is only driven L1CAM. We tracked the formation of trans-dimers in both scenarios. As shown in **Figure 4a**, the L1CAM system formed a higher number of trans-dimers compared to the NECTIN1 system. This indicates that L1CAM is more efficient at establishing the structural connections at cell-cell interface, because the trans-binding interface between L1CAM can coexist with its cis-interaction with the receptor. On the other hand, **Figure 4b** illustrates that the NECTIN1 system is more efficient to recruit FGFR1 receptors relative to the L1CAM system. Because the interaction between NECTIN1 and FGFR1 competes with NECTIN1 trans-dimerization, NECTIN1 is biased toward recruiting receptors, in contrast to the L1CAM system.

**Figure 4:**
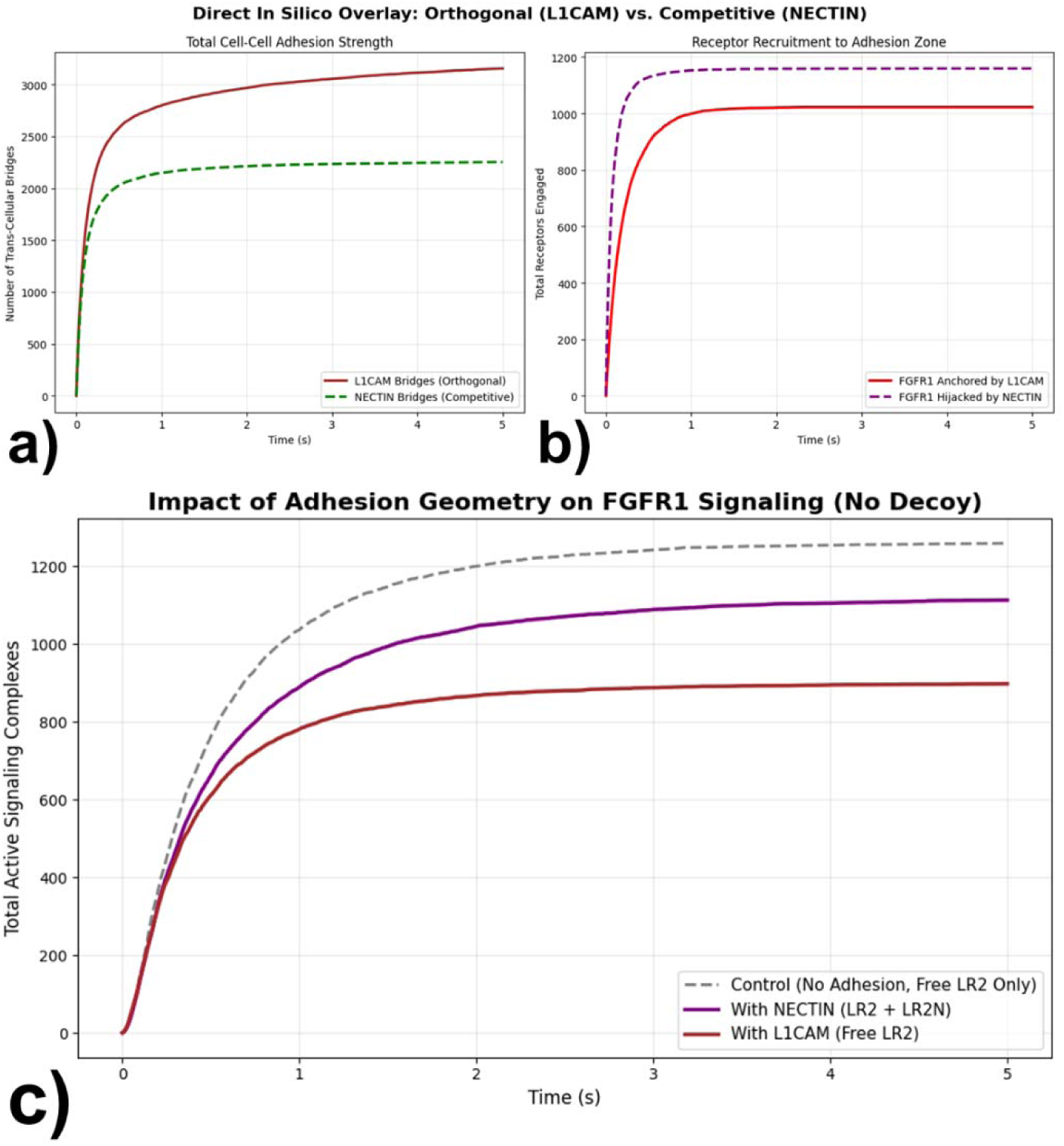
To understand how different cell adhesion molecules influence signaling, we compared two distinct systems: The first one is only driven by NECTIN1 and the second one is only driven L1CAM. We tracked the formation of trans-dimers in both scenarios (a). In contrast, the NECTIN1 system is more efficient to recruit FGFR1 receptors relative to the L1CAM system, as illustrated in (b). We further investigate how these adhesion systems affect the formation of the active ligand-receptor complexes needed for signaling. We compared these signaling outputs against a control system that lacks adhesion molecules. The results in (c) show that while L1CAM excels at forming structural connections between cells, this strong scaffolding effect comes at a cost to signaling efficiency. NECTIN1, while forming fewer adhesive interactions, recruits more receptors and allows a higher level of signaling to occur than L1CAM, though both adhesion systems still suppress signaling compared to a free-moving, non-adhesive membrane.

The next measure is how these adhesion systems affect the formation of the active ligand-receptor complexes needed for signaling. **Figure 4c** compares these signaling outputs against a control system that lacks adhesion molecules. Both the NECTIN1 and L1CAM systems resulted in fewer signaling complexes forming compared to the control system. This suggests that the process of being recruited and tethered to adhesion sites generally limits the receptors’ ability to form active signaling complexes. Furthermore, when comparing the two adhesion systems directly, the L1CAM system produced significantly fewer signaling complexes than the NECTIN1 system, because the NECTIN1-FGFR1 interactions do not overlap with FGFGR1 dimerization. In summary, while L1CAM excels at forming structural connections between cells, this strong scaffolding effect comes at a cost to signaling efficiency. NECTIN1, while forming fewer adhesive interactions, recruits more receptors and allows a higher level of signaling to occur than L1CAM, though both adhesion systems still suppress signaling compared to a free-moving, non-adhesive membrane.

The spatial information from the simulation is shown in **Figure 5**. The figure is organized into four grids, with each grid presenting a side-by-side comparison of specific molecular species at the Initial State (left, t = 0.0s) and the Final State (right, t = 5.0s). All heat maps represent a 15×15 voxels, where the color intensity indicates the number of molecules present in each voxel. **Figure 5a** shows the changes of free, unbound molecule before and after the simulation, with the FGF1 ligands presented in the upper panel and decoy receptor FGFRL1 in the lower panel. Relative to an initial random uniform distribution, the figure shows a more heterogeneous distribution formed at the end, due to their interactions with membrane receptors that are spatially localized on the plasma membrane together with other cell adhesion molecules. **Figure 5b** shows the changes in receptor-involved complexes before and after the simulation, with the active signaling complex formed between bivalent ligand and dimeric receptor presented in the upper panel, and the complex formed between FGFR1 and the decoy receptor FGFRL1 shown in the lower panel. These complexes are absent in the initial state but become highly abundant by the end of the simulation. Interestingly, the simulation reveals that they are not uniformly distributed across the two-dimensional membrane but instead form distinct clusters. Moreover, signaling complexes and decoy–receptor complexes tend to co-localize in close spatial proximity, likely because both arise from the same local pools of FGFR1 receptors and therefore preferentially form in membrane regions where receptor density is high.

**Figure 5:**
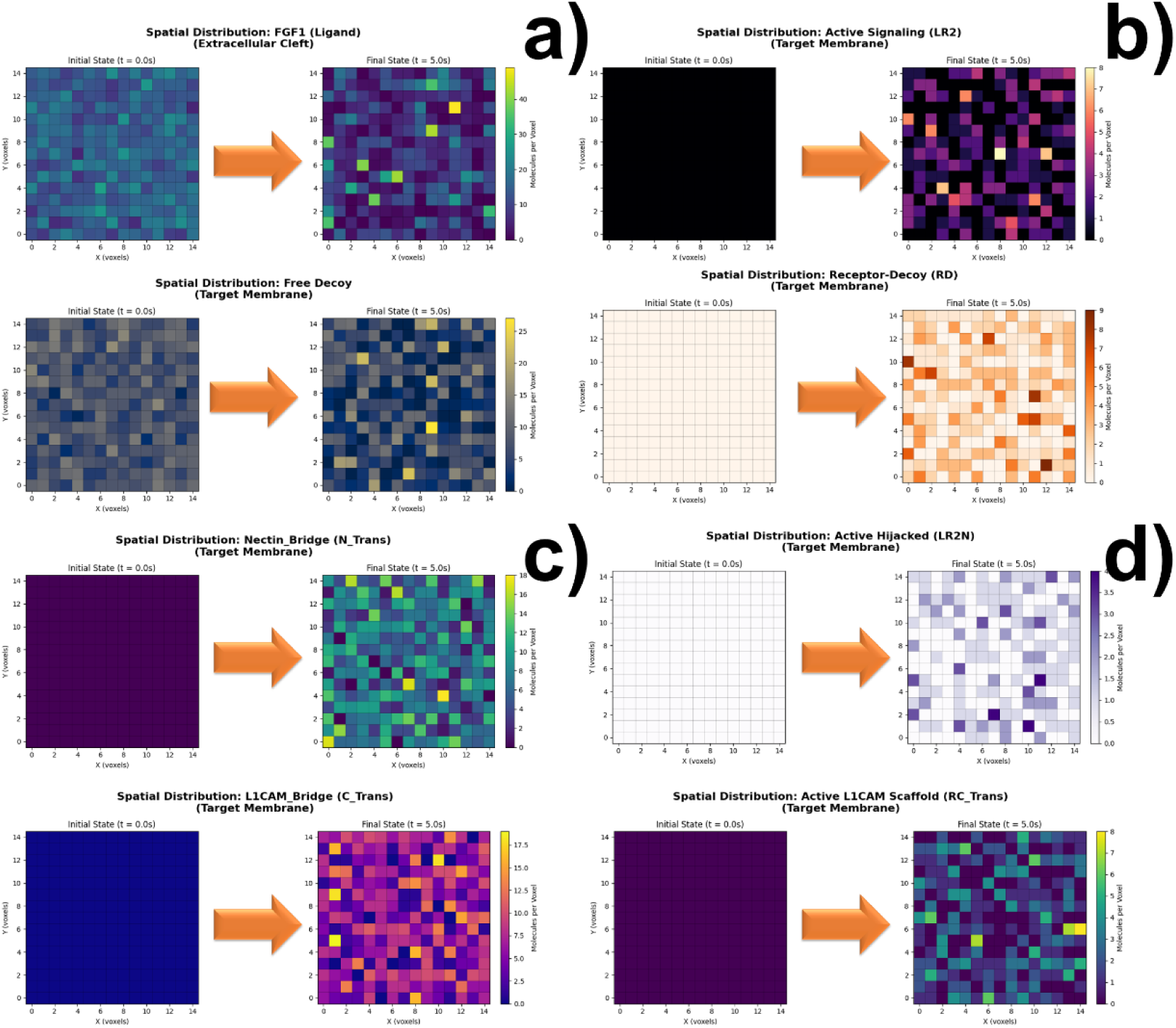
The spatial information from the simulation is organized into four grids, with each grid presenting a side-by-side comparison of specific molecular species at the Initial State (left, t = 0.0s) and the Final State (right, t = 5.0s). All heat maps represent a 15×15 voxels, where the color intensity indicates the number of molecules present in each voxel. The changes of free, unbound molecule before and after the simulation are shown in (a), with the FGF1 ligands presented in the upper panel and decoy receptor FGFRL1 in the lower panel. The changes in receptor-involved complexes before and after the simulation are shown in (b), with the active signaling complex formed between bivalent ligand and dimeric receptor presented in the upper panel, and the complex formed between FGFR1 and the decoy receptor FGFRL1 shown in the lower panel. The formation of trans-dimers along the simulation for NECTIN (upper panel) and L1CAM (lower panel) are shown in (c). Finally, the crosstalks between FGFR1 and its adhesion binding partners are shown in (d), with the growth of FGFR1-NECTIN1 interactions presented in the upper panel and growth of FGFR1-L1CAM interactions in the lower panel.

Figure 5c shows that formation of *trans*-dimers along the simulation for two cell adhesion molecules. The upper panel presents the growth of *trans*-interactions between NECTIN1, while the lower panel presents the growth of *trans*-interactions between L1CAM. Finally, Figure 5d shows the crosstalk between FGFR1 and its adhesion binding partners, with the growth of FGFR1-NECTIN1 interactions presented in the upper panel and growth of FGFR1-L1CAM interactions in the lower panel. Both *trans*-dimers and adhesion-signaling complexes show spatial heterogeneity. This probably is due to the fact that membrane receptors and adhesion molecules diffuse slowly on the two-dimensional membrane and become locally enriched through multivalent interactions, leading to the emergence of clustered micro-domains where signaling and adhesion complexes preferentially assemble.. Moreover, Figure 5c and **5d** provide visual evidence to Figure 4c that L1CAM creates a more stable and dense spatial “community” than Nectin. While Nectin’s competition between intercellular *trans*-interactions and *cis*-interactions with FGFR1 results in smaller, scattered hubs, L1CAM’s orthogonal geometry allows for the growth of larger, integrated signaling domains.

Together, these heat maps prove that cell signaling is not well-mixed, but forming discretized local hubs. Formation of these signaling or adhesive hubs suggests that the cell can trigger high-intensity local responses even when global molecular concentrations are low.

## Concluding Discussions

In conclusion, our multiscale computational framework demonstrates that intercellular communication is governed by strict physical and spatial constraints that cannot be captured by static interaction networks alone. By translating atomic-resolution structural interfaces into dynamic mesoscale simulations, we show that the cell–cell boundary is a highly partitioned and non–well-mixed environment. Adhesion molecules such as L1CAM and NECTIN1 function as key architectural organizers, restructuring the membrane into localized, high-intensity signaling hubs. At the same time, kinetic competition from decoy receptors plays a critical regulatory role. As illustrated by the D129A Kallmann syndrome variant, even subtle changes in binding affinity can redirect receptor fate and suppress essential downstream signaling. Together, these findings demonstrate that the structural geometry and spatial organization of multi-protein complexes are as fundamental as their biochemical concentrations in determining cellular responses. More broadly, this integrative framework provides a quantitative foundation for investigating the biophysics of cell–cell junctions and offers a promising strategy for identifying structurally informed therapeutic targets in diseases driven by dysregulated intercellular communication.

## Acknowledgement

This work was supported by the National Institutes of Health under Grant Numbers R01GM122804, and the United States – Israel Binational Science Foundation Project Number: 2023336. The work is also partially supported by a start-up grant from Albert Einstein College of Medicine. Computational support was provided by Albert Einstein College of Medicine High Performance Computing Center. Finally, we also greatly appreciate the general support from the Einstein 2030 Seed Fund.

## Author Contributions

Y.W. designed research; performed research; analyzed data; and wrote the paper.

